# Positivity effect in aging: Evidence for the primacy of positive responses to emotional ambiguity

**DOI:** 10.1101/2020.07.15.205096

**Authors:** Nathan M. Petro, Ruby Basyouni, Maital Neta

## Abstract

Older compared to younger adults show greater amygdala activity to positive emotions, and are more likely to interpret emotionally ambiguous stimuli (e.g., surprised faces) as positive. While some evidence suggests this positivity effect results from a relatively slow, top-down mechanism, others suggest it emerges from early, bottom-up processing. The amygdala is a key node in rapid, bottom-up processing and patterns of amygdala activity over time (e.g., habituation) can shed light on the mechanisms underlying the positivity effect. Younger and older adults passively viewed neutral and surprised faces in an MRI. Only in older adults, we found that amygdala habituation was associated with the tendency to interpret surprised faces as positive or negative (valence bias), where a more positive bias was associated with greater habituation. Interestingly, although a positive bias in younger adults was associated with slower reaction times, consistent with an initial negativity hypothesis in younger adults, older adults showed faster ratings of positivity. Together, we propose that there may be a switch to a primacy of positivity in aging.

## 1. Introduction

During normative adulthood, the transition into older age is accompanied by a decrease in the extent to which arousing, negative information impacts attention and memory (Mather, 2016). For instance, whereas in younger adults the processing of negative information is facilitated (Öhman et al., 2001) and interferes with attention toward competing neutral information (Müller et al., 2008), older adults show a reduction in this attention interference effect (Mather & Carstensen, 2003). Further, older compared to younger adults show less accurate memory recall of negative, but not positive, events (Charles et al., 2003). These age-related shifts away from negativity, customarily termed the “positivity effect” (Mather & Knight, 2005), are consistent with a general increase in reported emotional well-being among older adults (Charles, 2010).

Extensive neuroimaging work has examined the neural mechanisms underlying this positivity effect and has, for example, highlighted age-related changes in amygdala function. When viewing negative, but not positive, information, older compared to younger adults show less amygdala activation (Leclerc and Kensinger, 2011). Related work in older adults has linked the positivity effect to increased activation in the medial prefrontal cortex (mPFC; Dolcos et al., 2014; Leclerc & Kensinger, 2011; St. Jacques et al., 2010; van Reekum et al., 2018; Williams et al., 2006), suggesting that greater positivity may arise from a downregulation of amygdala activity via frontal cortical signals (Hariri et al., 2003). Taken together, this evidence supports the notion that the positivity effect emerges via top-down regulatory signals (Reed and Carstensen, 2012) which selectively control amygdala activity (Mather, 2016).

Another line of work has demonstrated that the positivity effect emerges during earlier time windows corresponding to early, low-level perceptual/attention processes, challenging the notion that the positivity effect results purely from top-down, emotion regulatory mechanisms. For instance, multiple studies leveraging the temporal resolution of electroencephalography have found, in response to negative facial expressions, there was no age-related difference in the relatively late brain potentials related to emotion processing (i.e., the P3 component; 350-450 ms; Vogel et al., 1998), but that the P1, a component reflective of early (70-130 ms) stages of perceptual processing (Hopfinger and Mangun, 1998), is diminished in older compared to younger adults (Houston et al., 2018; Mienaltowski et al., 2011). Using an emotion-induced blindness task, Kennedy and Mather (2019) found evidence that the positivity effect in older adults is present within the first 300 ms of stimulus presentation. Similarly, Gronchi and colleagues (2018) found that attention was attracted toward positive relative to negative faces at very brief (100 ms) presentations in older, but not younger, adults. These pieces of converging evidence suggest that the positivity effect emerges during early stages of perception, consistent with bottom-up processing.

The majority of work on the positivity effect relies on stimuli conveying clear positive or negative valence, and measures the age-related differences in a) attention shifts toward or away from competing positive and negative information, or b) the brain responses evoked by positive and negative information. More recently, work with dual-valence ambiguity (i.e., stimuli that could be validly interpreted as either positive or negative) have extended these findings by demonstrating the older adults have more positive interpretations of these stimuli than younger adults (Bucks et al., 2008; Neta and Tong, 2016; Shuster et al., 2017). Indeed, dual-valence ambiguity enables a measure of bias toward or away from positivity/negativity within a single item. For instance, whereas happy and angry expressions convey relatively clear positive and negative information, respectively, surprised expressions are ambiguous in that they signal both positive (e.g. an unexpected gift) and negative outcomes (e.g., witnessing a car crash). The increased positive ratings of surprised faces in older compared to younger adults (Neta & Tong, 2016; Shuster et al., 2017) suggests that, when the information within a single stimulus may convey multiple valid interpretations that are either positive or negative, older adults are more likely to ascribe positivity (i.e., *positive* valence bias).

The brain mechanisms underlying a positive valence bias have been explored, albeit exclusively in younger populations (i.e., children and younger adults). For instance, individuals with a more negative valence bias show increased amygdala and decreased vmPFC activity (Kim, Somerville, Johnstone, et al., 2003a; Petro et al., 2019). More recent evidence has shown that the same lateral frontal regions that are recruited during explicit emotion regulation are also recruited in response to surprised faces, but more so in individuals with a positive valence bias (Petro et al., 2018). These findings are consistent with other work that suggests that the initial response to dual-valence ambiguity (in young adults) is more negative, and that positivity may rely on a slower, regulatory process that overrides the initial negativity (Kaffenberger et al., 2010; Kim et al., 2003a; Neta et al., 2011, 2019; Neta & Tong, 2016; Neta & Whalen, 2010). These results might suggest that the positivity effect in older adults is the product of top-down signals which override the initial negativity in order to produce a more positive bias. However, given evidence that the positivity effect may emerge during temporally early time windows (Kennedy et al., 2019), we instead predict that there is a shift in older adulthood such that the initial response to dual-valence ambiguity is positive.

The goal of the present work is to explore the behavioral and neural mechanisms of the positive valence bias in older adults (compared to younger adults). Although some behavioral work in young adults has lent support for an initial negativity (e.g., longer reaction times for positive than negative trials, an initial attraction to the competing – negative – response when rating as positive; Brown et al., 2017; Neta et al., 2009), these approaches may be less compelling in aging. For example, behavioral responses such as reaction time and other motor responses show a general slowing in older age (Proctor et al., 2005). As such, we complemented our behavioral findings with neuroimaging measures to elucidate a more complete description of the mechanism supporting a positive valence bias in aging.

Specifically, the amygdala is considered a key node in the rapid, bottom-up processing of stimuli conveying biological relevance (LaBar and LeDoux, 1996; Sehlmeyer et al., 2009). With respect to facial expressions, the amygdala shows a robust response to negative (Johnstone et al., 2005; Whalen et al., 2001), positive (Costafreda et al., 2008; Sergerie et al., 2008), and even ambiguous (Kim, Somerville, Johnstone, et al., 2003a; Neta et al., 2013) expressions. Of note, although there is a robust response initially, the amygdala response to biological relevance habituates across repeated exposures (Bordi and LeDoux, 1992; Geissberger et al., 2020; Plichta et al., 2014), particularly when no further learning is required (Bordi et al., 1993; Breiter et al., 1996; Whalen et al., 1998). Interestingly, in the case of dual-valence ambiguity, while young adults show habituation to fear faces, there is relatively little change in the amygdala response to surprised faces (Whalen et al., 2009). The authors suggest that this lack of amygdala habituation for surprise may resemble slower extinction patterns because the ambiguity conveys uncertain outcomes which require further learning.

In the current study, we examined behavioral and neural (e.g., amygdala habituation) responses to surprised faces in older compared to younger adults. We predict that, consistent with previous studies (Neta & Tong, 2016; Shuster et al., 2017), older adults will show a more positive valence bias than younger adults. Further, given evidence that the positivity effect emerges early in time, we predict that older (but not younger) adults with a more positive bias will show greater amygdala habituation, suggesting that older adults perceive these expressions as more clearly positive (i.e., less in need of further learning). In other words, if the positive bias is a slower process that is preceded by an initial negativity (as seen in young adults), then the amygdala would putatively show sustained activity in response to an uncertain but potential threat. In contrast, if the positive response arises early on to signal safety, then no further learning would be required. In sum, we predict that, in contrast to what is seen in younger adults, a positive bias in older adults may be a faster response that is not characterized by an initial negativity.

## 2. Methods

### 2.1 Participants

Data were collected from 57 young (27 female, ages 17-30 years, mean(SD) age = 20.75(2.93)) and 52 older adults (36 female, ages 60-88 years, mean(SD) age = 69.92(6.83)) who reported having no history of neurological or psychiatric disorders, nor taking any psychotropic medication. The data from the sample of younger adults have been analyzed previously in Petro, Tong, Henley, and Neta (2018), but this previous analysis did not investigate effects related to amygdala habituation. During recruitment, older adults were administered the Modified Telephone Interview for Cognitive Status (Welsh et al., 1993); those with a score of 9/20 or higher on the recall portion of the interview and a total score of 24/39 or higher were invited to participate in the study. All recruitment and experiment protocols were approved by the University of Nebraska Committee for the Protection of Human Subjects in accordance with the Declaration of Helsinki. Each participant was given monetary compensation.

Among the recruited participants, 3 younger adults and 1 older adult failed to accurately rate clearly valenced facial expressions on at least 60% of the trials and thus were excluded from further analysis. This accuracy threshold has been used previously in young adult populations (Brown et al., 2017; Neta et al., 2009; Neta and Tong, 2016), and is a particularly important exclusionary criteria given the difficulty in discerning the specific interpretations of dual-valence ambiguity (i.e., valence bias) if stimuli with clear valence are not accurately rated. As such, 54 young (26 female, ages 17-30 years, mean(SD) age = 20.83(2.98)) and 51 older adults (36 female, ages 60-88 years, mean(SD) age = 69.94(6.90)) were included in the analysis of behavioral data.

In addition, 3 younger adults and 7 older adults did not complete the neuroimaging portion of the task (session 2, see below). The imaging data from 1 additional older adult were excluded due to technical failure during the session 2 task. Thus, 51 younger (25 female, ages 17-30 years, mean(SD) age = 20.73(2.93)) and 43 older adults (31 female, ages 60-88 years, mean(SD) age = 70.21(6.81)) were included in the analysis of MRI data.

### 2.2 Procedures

#### 2.2.1 Session 1: Valence Bias Assessment

See Figure 1 for an illustration of the experimental tasks. Session 1 comprised a behavioral testing session. All stimuli were presented on E-Prime software (Psychology Software Tools, Pittsburgh, PA, USA). To measure baseline valence bias, participants viewed images of happy, angry, and surprised facial expressions and rated (via keyboard press) each image as either positive or negative. The experimental design was taken from previous work (Neta et al., 2009). Stimuli included 34 discrete identities, 14 of which (7 females, ages 21-30 years) were drawn from the NimStim Set of Facial Expressions (Tottenham et al., 2009), and 20 from the Karolinska Directed Emotional Faces database (Goeleven et al., 2008). Each image was presented for 500 ms and separated by an interstimulus interval of 1500 ms. The images were presented across 2 blocks, each of which consisted of 24 images (6 angry, 6 happy, 12 surprise, per block) presented in a pseudorandom order, and blocks were counterbalanced between participants.

**Figure 1:**
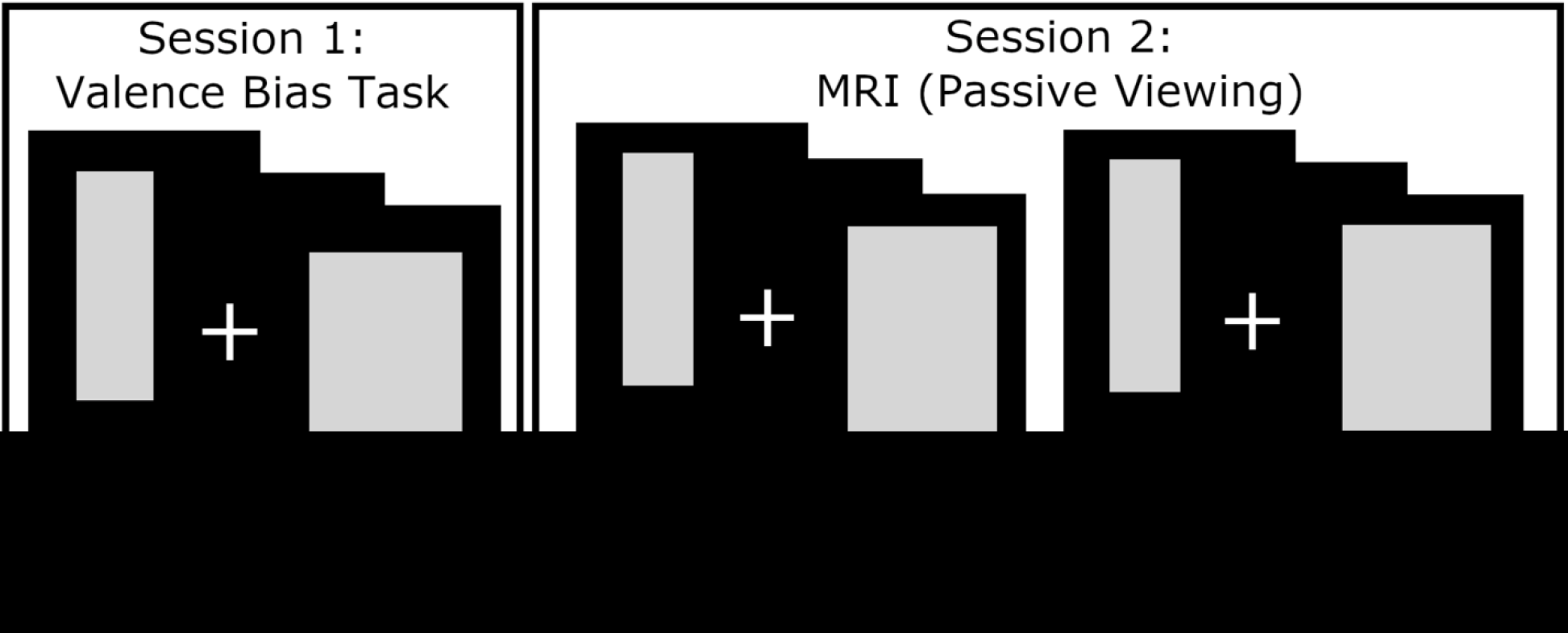
Depiction of procedure. In this illustration, gray boxes replace face stimuli as per bioRxiv requirements. In the valence bias task, participants viewed happy, angry, and surprised faces, and rated each image as either positive or negative. In the MRI, participants passively viewed a new set of faces (i.e., not seen in Session 1) during two runs with blocks of surprised and neutral faces. Faces depicted here are from the NimStim Set of Facial Expressions (Tottenham et al., 2009).

#### 2.2.2 Session 2: Magnetic resonance imaging (MRI)

Session 2 followed session 1 by approximately 7 days (Younger Adults: mean(SD) days = 7.84(2.09), range = 6-20 days; Older Adults: mean(SD) days = 7.09(0.87), range = 6-11 days). During the MRI scanning, participants freely viewed blocks of faces across four runs of blood-oxygen-level dependent (BOLD) imaging. For the younger adults, all stimuli during session 2 were presented using E-Prime software, whereas Experiment Builder (SR Research Ltd., 2015) was used to present stimuli to the older adult participants. The first two runs each consisted of 3 blocks of surprised faces and 3 blocks of neutral faces; the order these blocks were presented was pseudo-random. After these two runs, an additional two runs were completed in which fearful instead of surprised faces were presented, but the BOLD analysis of these runs is largely outside the scope of the current report (except for defining an amygdala region of interest; see below). Each individual block consisted of 32 faces (4 presentations of 8 unique identities), each presented for 200 ms and separated by a fixation cross for 300 ms, as in prior work (Kim & Whalen, 2009; Petro et al., 2018). Thus, the duration of each block was 16 seconds, and 14 seconds separated each face block during which a fixation cross was presented. Following these four runs, the participants completed two additional runs during which they completed an emotion regulation task, which is also outside the scope of the current report.

### 2.3 MRI acquisition and processing

#### 2.3.1 Scan parameters

The MRI images were collected in a Siemens 3T Skyra scanner using a 32-channel head coil at the University of Nebraska-Lincoln, Center for Brain, Biology & Behavior. The structural images were acquired using a T1-weighted MPRAGE sequence with the following parameters: TR = 2.2s, TE = 3.37 ms, slices = 192 interleaved, voxel size = 1.0 × 1.0 × 1.0 mm, matrix = 256 × 256, FOV = 256 mm, flip angle = 7 (degrees), total acquisition time = 5:07. BOLD images were collected while participants freely viewed the faces using an echo planar imaging (EPI) sequence with the following parameters: TR = 2.5 s, TE = 30 ms, slices 42 interleaved, voxel size = 2.5 × 2.5 × 3.0 mm, matrix = 88 × 88 mm, FOV = 220 mm, flip angle = 80 (degrees), total acquisition time = 3:24. The image slices were acquired parallel with the inter-commissural plane, and the volume positioned to cover the entire brain.

#### 2.3.2 MRI Preprocessing

Preprocessing of the imaging data was conducted using the Analysis of Functional Neuroimages (AFNI) suite of programs (Cox, 1996), and subsequent analysis of preprocessed imaging data was conducted in using both AFNI and MATLAB. The first four volumes of each run were discarded to allow for scanner stabilization. The BOLD timeseries, separately for each voxel, were first de-spiked by removing values with outlying data. Then, slice timing correction was accomplished by re-referencing each scan to the first slice. The slice time corrected volumes were then realigned to the minimum outlying image. All volumes were then aligned with the anatomical image, and then warped to the Talairach template atlas (Talairach and Tournoux, 1988) provided by AFNI. All functional volumes were then spatially smoothed using a 6mm^3^ full-width at half maximum kernel. The BOLD time-series, separately for each voxel, was normalized by dividing each time point by the average BOLD value across all time points and then multiplying all time points by 100. Any images containing movement exceeding 0.9 mm^3^, as calculated during spatial realignment, were censored frame-wise from further analysis.

### 2.4 Data Analysis

#### 2.4.1 Behavior

The valence bias score was calculated as the percent of negative ratings made for the surprised expressions out of the total number of ratings of surprise (i.e., excluding omissions). For example, a participant that rated all surprised faces as negative would be assigned a valence bias of 100%, but one that rated all surprised faces as positive would be assigned a valence bias of 0%.

Note that the ratings across expression conditions and across the two age samples were not normally distributed (all Shapiro-Wilkes *p*s < .05), thus non-parametric statistics are used for analyses of valence bias. For the moderation analyses, robust statistics were used in the regression.

To test whether valence bias was more positive in older compared to younger adults, the valence bias scores for each age group were submitted to a Wilcoxon signed-rank test. Further, given that previous work in younger adults has demonstrated that longer reaction times are associated with more positive ratings (Neta et al., 2009; Neta & Tong, 2016), we explored this relationship by submitting individual ratings of the surprised faces to a regression with three predictors: 1) reaction time of the rating, 2) age group, and 3) the interaction between reaction time and age group. The interaction term coefficient represented the moderating effect of age group on the relationship between valence bias and reaction time.

#### 2.4.2 Functional MRI - Amygdala BOLD Activation and Habituation

To identify amygdala voxels that were generally sensitive to face processing, the BOLD signal was submitted to a general linear model (GLM) containing regressors which modeled the stimulus onset and duration for each facial expression condition (surprised, fearful, neutral) separately. The regression matrix also contained 6 regressors modeling movement parameters (calculated during spatial realignment) and 2 regressors modeling polynomial trends to control for BOLD signal drifts. The beta values calculated from each task regressor were averaged together, separately at each voxel, and submitted to a one-sample t-test, yielding an estimate of the BOLD activation during the blocks of stimulus presentation. To identify amygdala activation, these t-values were passed through a cluster-forming (*p* < 0.001) and -extent (*k* = 23) threshold. This process revealed a cluster in both the right (*k* = 123, Talaraich (x, y, z) = 19, −6, −9) and left (*k* = 50, Talaraich (x, y, z) = −19, −6, −9) amygdala (Figure 2A). The dorsal position of these amygdalae regions is consistent with previous work showing that dorsal amygdalae are sensitive to stimuli conveying ambiguous valence and specifically to surprised and fearful facial expressions (Kim, Somerville, McLean, et al., 2003b; Whalen et al., 2001, 2009).

**Figure 2:**
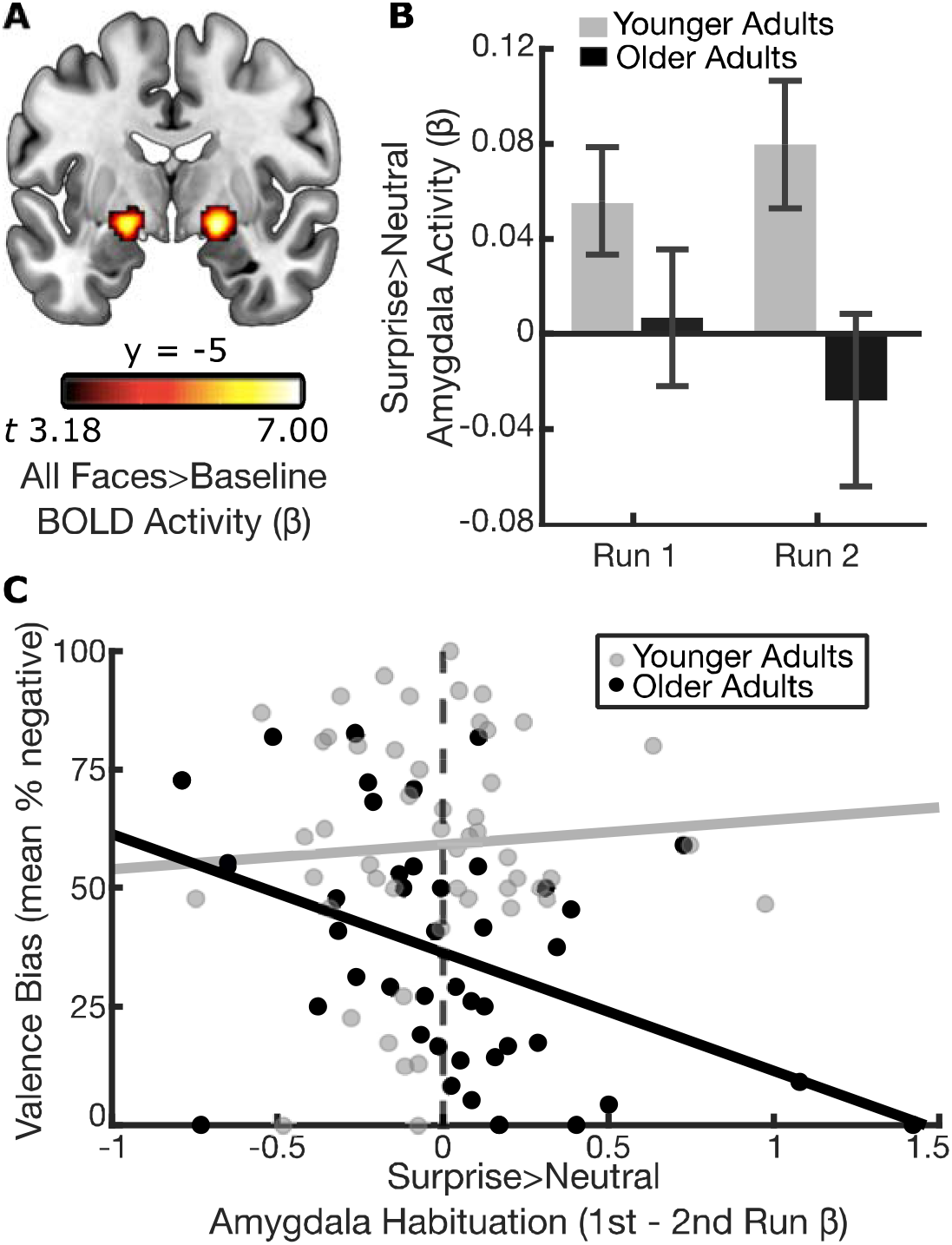
Relationship between amygdala habituation and valence bias in younger and older adults. **(A)** A seed region in the bilateral amygdalae was defined using the contrast of all facial expressions (surprise, neutral, fear) versus baseline (*p* < .001). **(B)** Surprise > neutral amygdala activity decreased from Run 1 to Run 2 for older (black bars) but not younger adults (gray bars). Error bars illustrate the between-subjects standard error. **(C)** Amygdala habituation was calculated as the difference in Run 1 > Run 2 for the surprise > neutral betas. Age group moderated the relationship between valence bias and amygdala habituation (*t*_90_ = 2.029, p = .045), such that greater amygdala habituation was related to more positive valence bias in older adults (in black; *t*_41_ = −2.991, p = .005) but not younger adults (in gray; *t*_49_ = 0.062, p = .951).

One goal of the present study was to analyze amygdala activity changes across runs of the experiment (i.e., habituation). To accomplish this goal, the BOLD time-series at each voxel was submitted to a GLM which contained regressors modeling the onset and duration of each stimulus block separately. Thus, a separate beta value was calculated for each stimulus block. The beta values within the bilateral amygdalae ROI were extracted and averaged across all voxels separately for each block. The block-by-block betas were first averaged together across all blocks for either facial expression condition (surprised and neutral), yielding an index of amygdala activity across the duration of the experiment for surprised and neutral expressions. In addition, to investigate the changes in amygdala activity between the two experimental runs, the beta values were averaged together for the 3 blocks within each experimental run (i.e., a single value per run), separately for each condition.

To determine whether amygdala activity evoked by expressions of surprise was related to valence bias, and if this relationship changed across age group, the surprise > neutral amygdala beta values, averaged across the entire experiment, were submitted to a regression with the outcome of valence bias and the predictors: 1) surprise > neutral amygdala betas, 2) age group, and 3) the interaction between surprise > neutral amygdala betas and age group. The interaction term coefficient represented the moderating effect of age group on the relationship between valence bias and amygdala activity. Further, in order to explore effects related to amygdala habituation, we also ran an additional regression replacing surprise > neutral amygdala beta values with surprise > neutral habituation values (Run 1 > Run 2 betas). In other words, these habituation betas were submitted to the same regression with predictors of valence bias, age group, and their interaction.

## 3. Results

### 3.1 Behavior

Both younger and older adults rated angry faces as negative (younger adults: mean(SD) % negative = 94.21(8.82); older adults: mean(SD) % negative = 90.99(9.23)) and happy faces as positive (younger adults: mean(SD) % negative = 6.43(8.91); older adults: mean(SD) % negative = 5.63(8.26)). In contrast, ratings of surprised faces showed more inter-subject variability for both younger (mean(SD) % negative = 58.69(23.89)) and older adults (mean(SD) % negative = 36.62(24.75)). For the purpose of the current study, the ratings of angry and happy faces, which convey information with clear valence, were used only as cut-offs for accurate performance. The ratings of surprised faces only were used to assess individual differences in valence bias. Consistent with previous work (Neta & Tong, 2016; Shuster et al., 2017), we found age-related differences in valence bias such that older adults rated surprised faces as more positive than younger adults (*z* = 4.13, p < .001).

For each age group, the reaction time of the ratings for surprised faces (younger adults: mean(SD) ms = 821.26(205.62); older adults: mean(SD) ms = 871.94(204.39)) were longer than for angry (younger adults: mean(SD) ms = 715.12(160.73); older adults: mean(SD) ms = 716.66(122.58)) and happy faces (younger adults: mean(SD) ms = 681.86(154.18); older adults: mean(SD) ms = 682.70(94.76)). Interestingly, there were no age-related differences in reaction times for rating surprised expressions (*z* = −0.96, *p* = .34).

The multiple regression revealed that age group moderated the relationship between valence bias and reaction time (*t*_101_ = 3.442, *p* < .001; Figure 3). Follow-up tests revealed that valence bias and reaction time were negatively related within the younger adults (*t*_52_ = −2.736, *p* = .009), consistent with previous work (Neta et al., 2009; Neta & Tong, 2016), but show a significant positive relationship within the older adults (*t*_49_ = 2.312, *p* = .025). In other words, younger adults showed a pattern whereby individuals with a negative bias rated surprised faces more quickly than those with a positive bias, but in older adults, as positive bias was associated with faster ratings.

**Figure 3:**
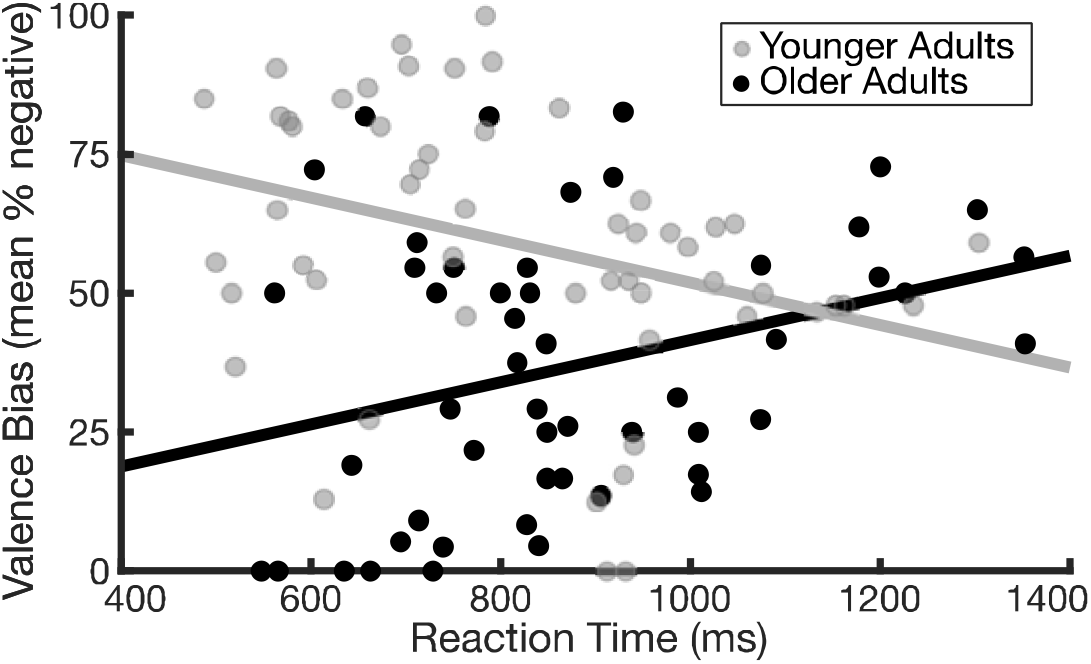
Relationship between valence bias and reaction time. Age group moderated the relationship between valence bias and reaction time (*t*_101_ = 3.442, *p* < .001). Longer reaction times were related to more positive valence bias in younger adults (in gray; *t*_52_ = −2.736, *p* = .009), consistent with previous work (Neta et al., 2009; Neta & Tong, 2016). Conversely, in older adults more positive valence bias was related to shorter reaction times (in black; *t*_49_ = 2.312, *p* = .025).

### 3.2 MRI

#### 3.2.1 Amygdala activity as a function of valence bias

When considering amygdala activity across all trials, the relationship between surprise > neutral amygdala activation was moderated by age group (*t*_90_ = 2.107, *p* = .038). Follow-up analyses revealed that older adults showed a negative relationship between surprise > neutral amygdala activity and valence bias (*t*_41_ = −2.982, *p* = .005), such that greater amygdala activity was related to more positive valence bias, whereas younger adults showed no relationship (*t*_49_ = −0.151, *p* = .881).

When considering patterns of habituation (Run 1 > Run 2) in the surprise > neutral amygdala activity, the relationship between surprise > neutral habituation and valence bias was moderated by age group (*t*_90_ = 2.029, *p* = .045; Figure 2B). Follow-up analyses revealed that older adults showed a negative relationship between surprise > neutral habituation and valence bias (*t*_41_ = −2.991, *p* = .005), such that greater amygdala habituation was related to a more positive bias. In contrast, younger adults showed no relationship between these variables (*t*_49_ = 0.062, *p* = .951). These differences between age group are not related to differences in the rate of habituation, since amygdala habituation did not differ between younger and older adults (t_92_=−0.776, *p* = 0.440).

To probe this relationship between amygdala habituation and valence bias in older adults, the amygdala activation betas for both surprise and neutral blocks separately, for both Run 1 and Run 2 separately, were submitted to a bivariate regression with valence bias, resulting in a total of 4 regression analyses (Figure S1). In other words, there was a regression for 1) surprise-related activity in Run 1 and 2) in Run 2, and for 3) neutral-related activity in Run 1 and 4) in Run 2. During Run 1, surprise-related activity was negatively related to valence bias (*t*_41_ = −2.579, *p* = .014), such that greater amygdala activity predicted a *more* positive valence bias. In contrast, neutral-related activity in Run 1 was positively related to valence bias (*t*_41_ = 2.125, *p* = .040; i.e., greater amygdala activity predicted a more negative bias). During Run 2, neither surprise-nor neutral-related activity were related to valence bias (surprise: *t*_41_ = −0.662, *p* = .511; neutral: *t*_41_ = −1.520, *p* = .136).

Finally, given that there was some variability in the rate of habituation in older adults including a subset of individuals showing an enhanced amygdala response over time, we further explored these habituation effects. As illustrated in Figure 4, those with negative values (black dots, n = 21; mean(SD) = −0.258(0.235)) show a repetition enhancement, while positive values (white dots, n = 22; mean(SD) = 0.314(.352)) represent repetition suppression, or habituation.

**Figure 4:**
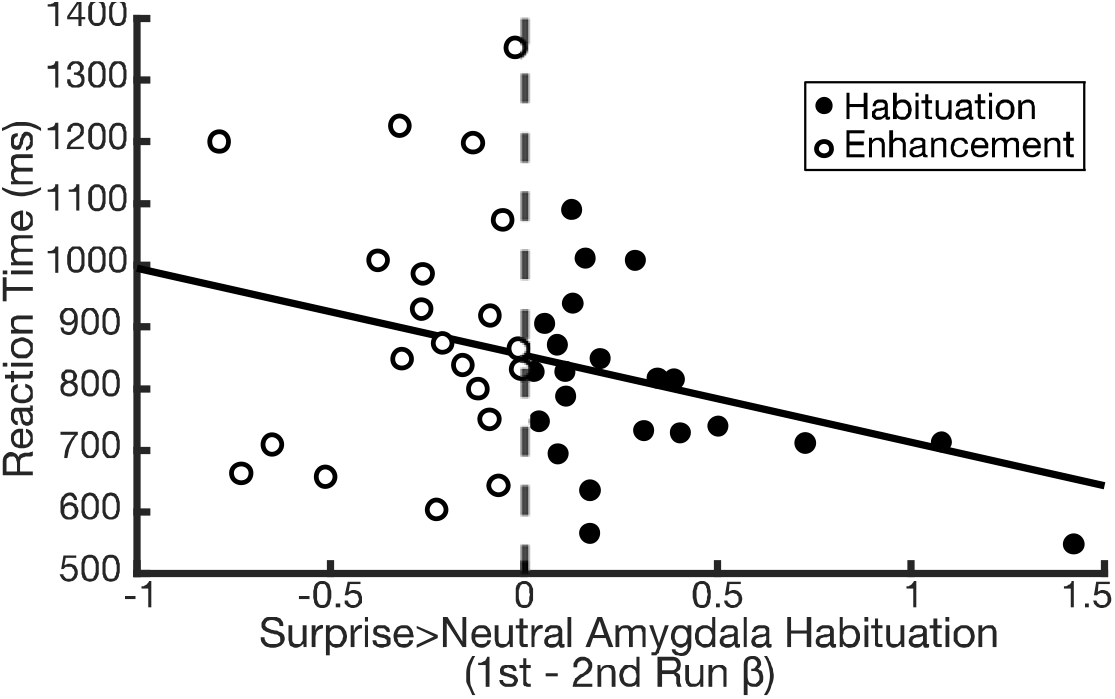
Variability in amygdala habituation and reaction time in older adults. From the 1^st^ to the 2^nd^ run, 21 individuals showed increased amygdala activity over time (white filled dots; mean(SD) = −0.258(0.235)) and 22 showed decreased activity across runs (black filled dots; mean(SD) = 0.314(.352)). Among this variability in amygdala change, those who showed greater habituation of amygdala activity had shorter reaction times (*r*_41_ = −.277, *p* = .073).

One potential explanation for these differences in older adults might be related to cognitive function. To test this, we operationalized cognitive function using fast reaction times, such that faster reaction times represented relatively better cognitive function. We then compared reaction time with changes in amygdala activity. Each participant’s average reaction time for ratings of surprised face ratings was calculated, and then submitted to a Spearman’s rank correlation with amygdala habituation. This test revealed that, if anything, greater habituation was weakly associated with faster reaction times (Figure 4; *r*_41_ = −.277, *p* = .073).

## 4. Discussion

Older compared to younger adults rated the expressions of surprise as more positive, replicating previous work (Neta and Tong, 2016; Shuster et al., 2017) and broadly consistent with the positivity effect in aging (Mather, 2016; Mather & Knight, 2005). Importantly, the observed relationship between valence bias and both 1) reaction times and 2) amygdala habituation suggests that this age-related shift toward more positivity is accompanied by a shift away from an initial negativity. In other words, older adults show a faster or more default positivity, rather than negativity, in response to dual-valence ambiguity.

Indeed, the notion that the positive valence bias in aging results from a default response that is more positive is supported by the age-related differences in reaction times. Specifically, in younger adults, more positive ratings of surprised faces were associated with slower reaction times (Neta et al., 2009), and an instruction to deliberate is sufficient for promoting greater positivity (Neta and Tong, 2016). This effect replicates extant literature and is thought to reflect additional, putatively top-down, regulatory signals (Kaffenberger et al., 2010; Neta et al., 2011; Neta and Tong, 2016; Petro et al., 2018). However, the opposite pattern was found in older adults, in which positive ratings were related to faster reaction times. This age-related difference in the relationship between valence bias and reaction time suggests that while younger adults appear to rely on regulatory processes (characterized by longer reaction times) for the positive evaluation of ambiguous emotional content, a different process (characterized by shorter reaction times) may drive positive evaluations in older adults.

Moving to the neuroimaging findings, first, we replicated previous work showing no evidence for amygdala habituation in response to surprised faces (Whalen et al., 2009). Further, as predicted, older (but not younger) adults showed a relationship between habituation and valence bias, such that those with a more positive valence bias showed greater habituation. Interestingly, patterns of persistent (as opposed to habituating) amygdala activity across stimulus presentations reflect a brain mechanism which maintains, across repeated exposures, the processing of biologically relevant information to promote learning (Davis et al., 2016; Herry et al., 2007; Whalen, 2007). In the context of the extant literature on amygdala habituation, it could be that younger adults perceive the potential (but ambiguous) negativity in response to surprised faces and thus require further learning, whereas the older adults perceive the potential positivity and thus no further learning is required.

To elaborate, the change in amygdala activity across runs showed wide variability in the older adults. For example, while some individuals showed habituation, others showed response enhancement. We found that the former group was more likely to have a positive valence bias, while the latter group was more likely to have a negative valence bias. Again, one possible explanation is that the amygdala increase over time is a characteristic of the latter group perceiving potential negativity or threat in response to the surprised faces. And, given there is some uncertainty about the presence of this threat, the amygdala stays active to promote further learning, consistent with the pattern previously observed in younger adults (Davis et al., 2016). Conversely, the individuals with a more positive valence bias appear to be more likely to perceive positivity in response to the surprised faces, and thus render the faces “safe” or not in need of further learning. This speculative interpretation is supported by age-related differences in the relationship between valence bias and reaction time. Specifically, although younger adults are faster when rating surprised faces as negative than positive (see also Neta et al., 2009), older adults are faster when rating the same faces as positive.

Another possibility is that the individual differences in amygdala change across runs are related to cognitive function. Indeed, cognitive function generally declines with age (Salthouse et al., 2003), and it could be that the amygdala habituation is associated with greater cognitive deficit. Although no explicit measure of cognitive function was collected in the current study, we explored the possibility that slower reaction times might be a useful proxy for cognitive decline. Interestingly, older adults with greater decreases in amygdala activity showed faster (not slower) reaction times. Thus, these results suggest that amygdala habituation may be putatively related to *better* cognitive function, consistent with work showing that those with relatively good cognitive function tend show a stronger positivity effect (Mather & Knight, 2005). Future studies will benefit from adding an explicit measure of cognitive ability in order to further explore this pattern of results.

The surprise and neutral amygdala activity evaluated separately across both runs suggests that this habituation effect in older adults with a more positive valence bias is characterized in part by an initial amygdala increase toward surprised faces. The finding that increased amygdala activity is related to more positivity is consistent with prior work which found that older adults show increased amygdala activity to positive relative to neutral and negative information (Kehoe et al., 2013; Leclerc & Kensinger, 2011; Mather et al., 2004), particularly for relatively low-arousing stimuli (Dolcos et al., 2014). In terms of the habituation effect, one straightforward explanation for this pattern of results is that the amygdala activity increases initially, and the greater increase allows for a greater change (habituation) in the second run. In contrast, the individuals with a more negative bias show no change in amygdala activity or show an increase in the second run (see more on this below). Despite the variability in habituation across individuals, we suggest that the pattern of habituation in the subset of older adults with a positive bias is novel, given that previous work has only evidence for sustained amygdala activity in response to surprise (as opposed to fearful) expressions (Whalen et al., 2009).

Notably, the pattern of amygdala activity across runs in older adults is specific to surprised faces and not general to neutral faces. Indeed, a previous literature has demonstrated that older adults are less efficient at differentiating facial features (i.e., de-differentiation) relating to identity (Bartlett et al., 1989; Goh et al., 2010) and emotion (Orgeta and Phillips, 2007). Thus, in the context of the current results, one possibility is that greater habituation of amygdala activity and more positive valence bias are associated with greater levels of de-differentiation. In other words, the amygdala may habituate more in older adults who are less able to discern the differences between each face or identify their expressions. However, in the current results the relationship between amygdala habituation and valence bias was observed uniquely for surprised (but not neutral) faces, indicating that the amygdala response to these different expressions is indeed sensitive to the content of the expressions.

The interpretation of the current results, that positivity in older adults constitutes an early and default response, is qualified by limitations of the methodology used. Foremost, the measurements of amygdala activity depend on the sluggish BOLD signal which is unable to dissociate temporally early from late processes. Other studies using EEG or rapid stimulus presentations have demonstrated that a bias towards positive information in older age occurs during relatively early time windows (Gronchi et al., 2018; Houston et al., 2018; Kennedy et al., 2019; Mienaltowski et al., 2011). Future work may utilize these temporally precise methods to test our prediction that, in older age, positive compared to negative interpretations of dual-valence ambiguity involve relatively early processes. Similarly, eye-tracking measures are well suited to track attention processes during these fleeting, early time-windows.

Finally, previous work has found that younger adults with a more negative valence bias show greater surprise-related amygdala activity than those with a positive bias (Kim, Somerville, Johnstone, et al., 2003a). However, in our data, there was no relationship between valence bias and amygdala activity in younger adults. Methodological differences may account for this inconsistency. For instance, Kim and colleagues (2003b) used a whole-brain correlation conducted on the surprise > baseline beta values to identify a cluster in the more ventral region of the amygdala. Further, valence bias was measured using only a single trial immediately following the MRI session. In the current study, conversely, the amygdala cluster was defined as voxels showing activation to all faces, and its voxels where relatively dorsal. Further, we calculated valence bias throughout an entire behavioral task (consisting of approximately 24 trials), conducted approximately a week before the MRI session. This discrepancy in the relationship between valence bias and amygdala activity in young adulthood may be explored in future work.

### 4.1 Conclusions

To summarize, older, but not younger, adults showed amygdala habituation to surprised faces. The magnitude of habituation was associated with a more positive valence bias and shorter reaction times when rating the valence of surprised faces. These results suggest that, whereas a positive valence bias in younger adults putatively relies on an additional regulatory mechanism, older adults show evidence for a primacy of positivity in response to dual-valence ambiguity.

## Acknowledgements

This work was supported by the National Institutes of Health (NIMH111640; PI: Neta), the National Science Foundation (CAREER Award; PI: Neta), and by Nebraska Tobacco Settlement Biomedical Research Enhancement Funds. We thank Elizabeth Kensinger for helpful comments on the manuscript. We also thank Kayla Clark, Daniel J. Henley, and Tien T. Tong for assistance in data collection and management.

