## Supplemental Figure 1 for "Positivity effect in aging: Evidence for the primacy of positive responses to emotional ambiguity"

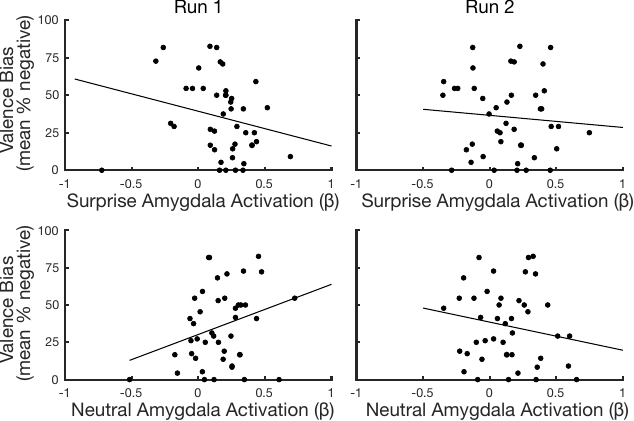

**Figure S1:** **Relationship between valence bias and amygdala activity across either run and condition.** During Run 1 (left column), more positive valence bias was related to greater amygdala activity to surprised faces (top left; *t*_41_ = -2.579, p = .014), but lower amygdala activity to neutral faces (bottom left; *t*_41_ = 2.125, p = .040). During Run 2 (right column), valence bias was not related to amygdala activity evoked by either surprised (top right; *t*_41_ = -0.662, p = .511) or neutral faces (bottom right; *t*_41_ = -1.520, p = .136).
